# Localization accuracy in single-molecule fluorescence microscopy

**DOI:** 10.64898/2026.10.14.759109

**Authors:** Katarina B. Chapman, Roman Schmidt, Johann Engelhardt, Stefan W. Hell

## Abstract

The two prevalent methods to determine the position of individual fluorophores in the focal plane of a microscope rely either on the centroid of the fluorescence pattern produced by the fluorophore on a camera or on targeting the fluorophore’s coordinate with a spatially controlled illumination minimum, as realized in MINFLUX. In both methods, (sub)nanometer localization precisions have been reached, but little is known about their accuracy, i.e., the deviation of the obtained coordinate from its ground truth. Here we report a theoretical and experimental comparison of the accuracy provided by these two strategies. We quantify substantial systematic errors in camera-based localization due to the interplay of two difficult-to-control parameters: the orientation of the fluorophore in space and its axial deviation from the focal plane (defocus). For example, within an axial range of ±120 nm around the focus, the median distance between two fluorophores on a nanobody measured by camera-based localization varies by 26.5 nm. Under the same conditions, a commercial MINFLUX system yields a median variation in distance of 1.4 nm, indicating an improvement in accuracy by an order of magnitude. The accuracy of MINFLUX localization can be further improved to (sub-)ångström levels by reducing the average distance between the coordinates targeted by the illumination minimum and the fluorophore. As it delivers unrivaled precision and accuracy for mapping fluorophores and their distances, we consider coordinate-targeted localization essential for studying labeled cells at the sub-10 nm level and for exploring structural and conformational changes of labeled biomolecules.

---

Determining the position of individual fluorophores is vital in superresolution fluorescence microscopy^1,2^. Currently, two fundamentally different methods exist for localizing a single fluorescent molecule^3,4^. In camera-based localization, as used in PALM^5,6^, STORM^7^, and PAINT^8^, the molecular position is inferred from the center of the diffraction pattern that the fluorescent molecule produces on a camera. In this case, the localization precision scales with the fluorescence wavelength and inversely with the square root of the number of detected photons *N*. Camera-based localization thus relies entirely on the light emitted by the molecule itself. Coordinate-targeted localization methods, on the other hand, delegate the need for many photons for defining a coordinate to the illumination beam. Concretely, in the concepts called MINFLUX^9^ and MINSTED^10,11^, the molecule is localized by relating its unknown position to a known coordinate in the sample that is established by the intensity zero of a typically doughnut-shaped illumination beam. The intensity zero is targeted as close as possible to the fluorophore. Then, the fluorophore is localized by referencing its coordinate to the controlled position of the zero. Contrary to its camera-based counterpart, in coordinate-targeted localization the attainable precision is not dominated by the wavelength but scales with the diameter *L* describing the targeted area around the fluorophore. The smaller the *L*, the higher the precision with which the molecule can be localized with a given number *N* of detected photons^9^.

Both these concepts have achieved localization precisions down to the ångström level. Camera-based localization attained this feat by ever increasing *N* to the tens or hundreds of thousands, and even millions^12–15^, whereas coordinate-targeted methods achieved single-digit ångström precisions^11,16–18^ with orders of magnitude fewer detected photons by minimizing the distance between the zero and the molecule, *L*. Nonetheless, compared to the localization precision, which describes the spread of repeated position estimates, much less is known about the attainable accuracy, that is, how closely the estimated position corresponds to the true position of the molecule.

Systematic errors in camera-based localization were recognized and studied early on^19–22^. These errors ultimately stem from the fact that fluorescence emission is a molecular dipole transition that, in conjunction with even a slight defocus (*z* ≠ 0), makes the fluorescence pattern on a camera heavily depend on the emitter’s orientation in space^23,24^. For rotationally restricted molecules with *z* ≠ 0, almost any dipole orientation displaces the center of the emission pattern from the coordinate that mirrors the molecular position on the camera^19–22^. Solutions have been given to correct these errors^25–28^, but these strategies come with their own limitations and have not become widely applied. It has been argued that MINFLUX is substantially less susceptible to such biases because its spatial reference coordinate is defined by a controlled illumination rather than a varying emission pattern^9^. However, this expectation has so far been based on reasoning^29^ and theory only^29^, with no direct experimental proof. Here, we provide rigorous theoretical and experimental evidence that MINFLUX delivers substantially higher localization accuracy than popular camera-based localization. Coordinate-targeted localization, as implemented in MINFLUX and also MINSTED, thus becomes critical for obtaining accurate molecular coordinates of individual fluorophores at (sub)nanometer length scales.

## Results

### MINFLUX is intrinsically less susceptible to dipole-induced errors than emission-based camera localization

The difference between the two optical localization strategies is fundamental: the camera-based scheme demands from the fluorophore to yield all the photons needed to establish its coordinate (Fig. 1a), whereas its coordinate-targeted counterpart injects a photon pattern provided by the instrument (Fig. 1b). On the molecular level, a fluorophore interacts with light through its electric transition dipole moment ***µ***(*θ, φ*), where *θ* is its inclination angle from the optical axis and *φ* is the azimuthal angle in the sample plane (Fig. 1a,b). The component of the excitation field parallel to the dipole drives the transition^30,31^, so the excitation rate is proportional to |***E*** · ***µ***(*θ, φ*)|^2^. Likewise, the excited dipole emits fluorescence in a toroidal pattern with no emission along the axis and maximum emission perpendicular to it. The objective lens collects the emitted light so that its field yields a two-dimensional diffraction pattern, essentially forming a spot in the image plane^32,33^. In camera-based methods, the fluorophore’s position is derived from this fluorescence spot by centroid calculation, Gaussian fitting or related methods (Fig. 1a). The underlying assumption is that, when back-projected to the sample plane, the calculated centroid or fitted peak reflects the actual molecular position.

**Fig 1.**
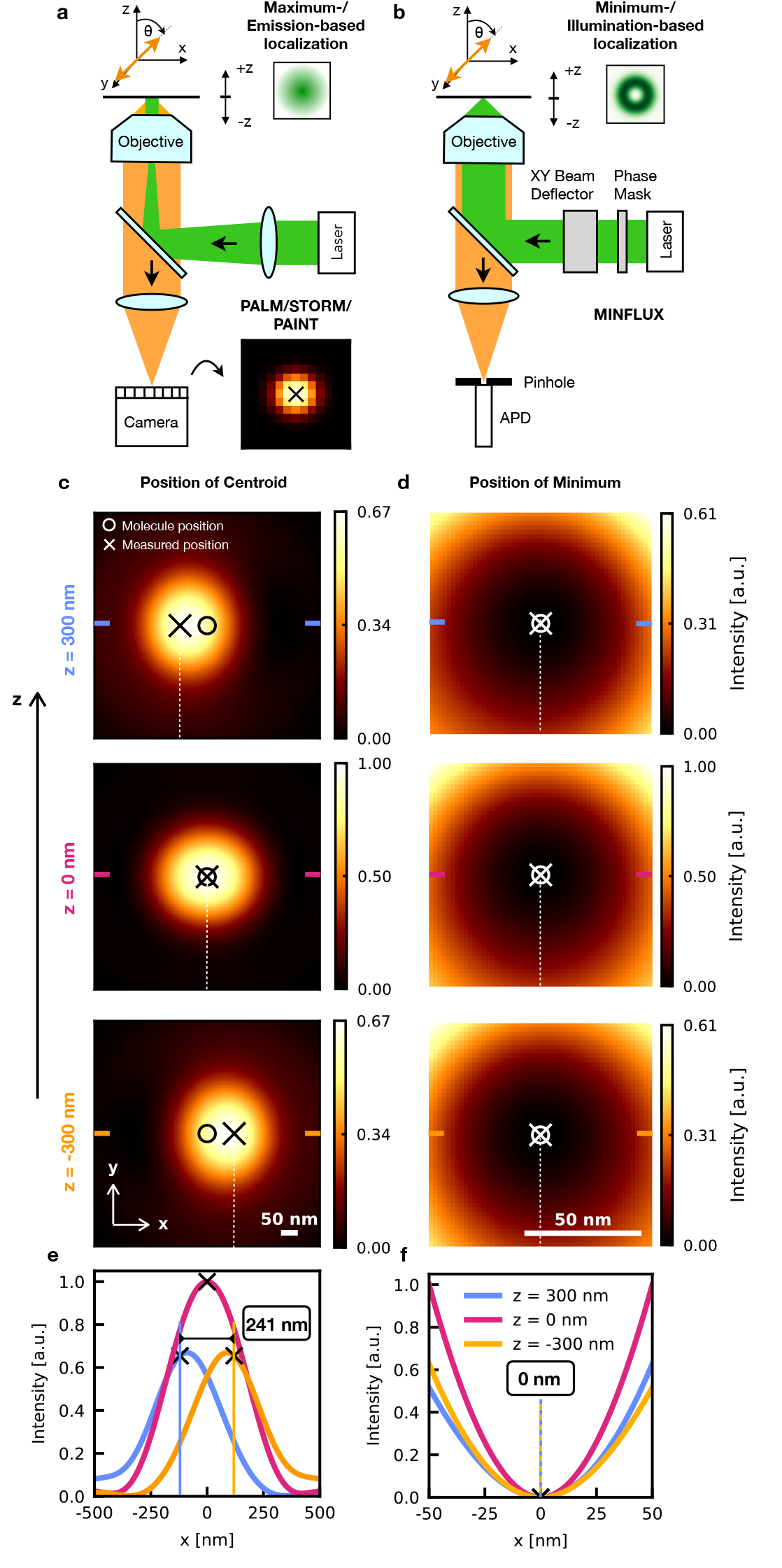
Emission-image-based and excitation-minimum-based localization of a spatially oriented fluorophore. **a**,**b**, Schematics of camera-based localization as used in PALM/STORM and PAINT (**a**) and MINFLUX localization using a confocal setup with a doughnut-shaped excitation beam featuring a central intensity zero (**b**). Camera-based methods estimate the molecular position from the diffraction pattern of fluorescence recorded on a camera by associating it with the centroid of the pattern. MINFLUX estimates the position from photon counts recorded with the excitation minimum placed at known positions around the molecule, by associating it with the coordinate where molecular emission is expected to vanish. **c**,**d**, Simulated camera images of single molecule fluorescence (**c**) and scanned fluorescence intensity distributions recorded by the confocal detector (**d**) for a fixed dipole inclined by θ = 45° at defocus z = −300, 0 and +300 nm (NA = 1.4, index-matched sample with n = 1.518; emission/excitation wavelengths of 670/640 nm). Circles mark the true molecular position; crosses mark the estimated position (intensity-weighted centroid within ±1 µm in **c**; fluorescence minimum in **d**). **e**,**f**, Horizontal profiles through the distributions in **c** and **d**. The camera-image centroid shifts by 241 nm between defocus z = −300 and +300 nm (**e**), whereas the MINFLUX excitation minimum and hence also the fluorescence minimum always remain exactly at x = 0 (**f**).

Unfortunately, the fluorophore’s unknown orientation, rotational freedom and axial displacement (*z* ≠ 0) all influence the pattern on the camera. For an immobilized or rotationally restricted molecule, even a small axial defocus (*z* ≠ 0) induces notable changes on the emission wavefront, causing the back-projected center of the pattern to no longer correspond to the actual position of the fluorophore. In fact, asymmetric phase shifts in high-aperture imaging move the pattern laterally even though the molecule itself has not moved. For a dipole inclined by 45°, translating the fluorophore from *z* = -300 nm to +300 nm, or vice versa, shifts the centroid of the fluorescence pattern by 241 nm in the focal plane (Fig. 1c,e). This applies to camera localization with a numerical aperture (NA) of 1.4, a fluorescence wavelength of 670 nm and an index-matched sample (*n* = 1.518). The same conditions were also applied when calculating the performance of MINFLUX localization, except that we considered the relevant excitation wavelength of 640 nm.

In a confocal MINFLUX setup with a doughnut-shaped excitation beam with a central intensity zero, the fluorescence image resulting from scanning the minimal intensity center of the beam across the fluorophore behaves differently: the central point of vanishing fluorescence does not move at all (Fig. 1d,f). As a matter of fact, the position of the fluorescence minimum depends neither on the orientation of the molecule nor on its defocus. The reason is that a true zero of the excitation field (***E*** = 0) implies |***E*** · ***µ***(*θ, φ*)|^2^ = 0 for any dipole orientation. This is the defining accuracy advantage of coordinate-targeting. Since the coordinate of the zero is injected by the laser beam, the position at which the signal disappears is a property of the illumination light field alone and thus independent of *θ, φ* and *z*.

Practical MINFLUX localization does not literally impose a match of the position of the excitation zero with that of the fluorophore. Instead, it estimates the coordinate where the zero would cause the fluorophore to go dark. This estimated coordinate is then regarded as the position of the molecule. Since MINFLUX localization still draws on fluorescence signal for localization, the orientation of the molecule plays a role, albeit to a lesser extent than in the camera case. For an inclined dipole, orientation-dependent coupling to the vectorial excitation field makes the fluorescence pattern around the zero slightly asymmetric, as indicated by the yellow and blue curves in Fig. 1f. In contrast, on a camera, a *z* = 50 nm defocus of a fluorophore inclined by *θ* = 45° displaces the calculated centroid laterally by 20.5 nm (Fig. 2a, upper panel). MINFLUX localization of the same dipole, using the six-point estimator implemented in the commercial instrument used further in the experimental part of our study (Abberior Instruments GmbH, Göttingen, Germany)^34^ and *L* = 10 nm is predicted to yield a systematic error of only 0.53 Å (Fig. 2a, lower panel). This minor sub-ångström error stems from the fact that high-aperture focusing of a circularly polarized field generates not only focal plane components *E*_*x*_ and *E*_*y*_, but also a longitudinal component *E*_*z*_ featuring a different phase behavior. As an inclined molecule couples to all field components, its excitation rate is co-determined by their phase differences making *E*_*z*_ and *E*_*x,y*_ add constructively on one side of the zero and destructively on the other.

**Fig 2.**
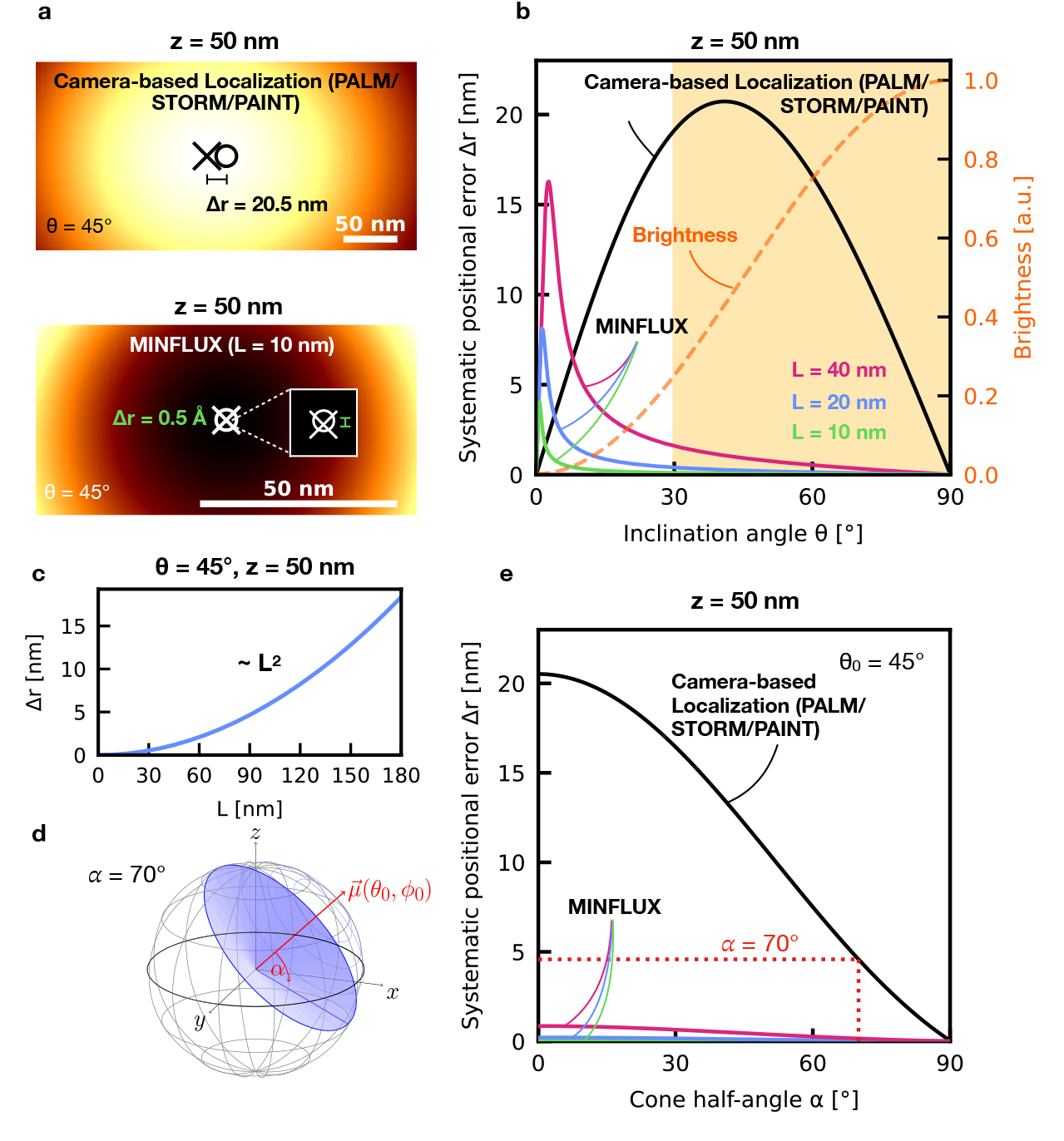
Simulated orientation-dependent systematic positional (accuracy) errors for fixed and partially rotating dipoles. **a**, Simulated camera intensity distribution (top) and MINFLUX fluorescence intensity distribution (bottom) for a fixed dipole with θ = 45°, ϕ = 0° and z = 50 nm. Circles and crosses mark the true and estimated positions, respectively. Simulation parameters as in Fig. 1; MINFLUX positions were calculated from the fluorescence intensities at six equally spaced probe positions on a circle of diameter L using the commercial instrument’s estimator. The systematic positional error is Δr = 20.5 nm for camera localization and 0.05 nm for MINFLUX with L = 10 nm. **b**, Δr as a function of θ for camera localization and MINFLUX with L = 40 nm (magenta), 20 nm (blue) and 10 nm (green). The orange dashed curve shows the normalized fluorophore brightness; shading denotes orientations satisfying a brightness B(θ) > B_max_/4. **c**, Δr as a function of L for θ = 45° and z = 50 nm, displaying the quadratic dependence on L. **d**, Cone model of rotational mobility, defined by mean orientation (θ_0_, ϕ_0_) and half-angle α. **e**, Δr as a function of α for θ_0_ = 45°. The red dotted line marks α = 70°, corresponding to the cone illustrated in **d**.

### Residual MINFLUX systematic error decreases quadratically with probing diameter *L*

For MINFLUX, the distance *Δr* between the estimated and true fluorophore positions depends to a minor extent on the dipole orientation (Fig. 2b) and probing diameter *L* (Fig. 2c), but hardly on defocus. Defocus changes the estimated *x*- and *y*-coordinates: the bias vector changes direction, and the estimated position moves around the fixed excitation zero, while its radial magnitude *Δr* remains approximately constant. Camera-based localization behaves differently, as *Δr* increases approximately linearly with |*z*| over the investigated range. This is why camera-based localization entails large displacements (Fig. 1e) at large defocus. However, even at a minor defocus of *z* = 50 nm, camera-based systematic errors can reach tens of nanometers also for bright, detectable fluorophore orientations (Fig. 2b).

On the other hand, for MINFLUX, the largest possible errors occur only for fluorophores with inclination angles *θ* < 10°, i.e., those that are essentially perpendicular to the focal plane, hence dim and practically irrelevant (Fig. 2b). This inclination dependence stems from the contribution of both the transverse and longitudinal excitation fields. The leading asymmetric contribution scales as sin *θ* cos *θ*, whereas the leading symmetric excitation signal scales as sin^2^ *θ*; their ratio therefore increases approximately as |cot *θ*|. The systematic error decreases rapidly for fluorophores with larger inclination angles that are inherently brighter. Its leading contribution vanishes for an in-plane orientation, *θ* = 90°, for which the axial dipole component is zero. In an ideal rotationally symmetric system, the azimuthal angle *φ* determines only the direction of the bias vector and has no effect on its magnitude. The same applies to a change of the arrangement of the six targeted coordinates in the focal plane that surround the fluorophore to be localized.

An important outcome is that, in all relevant cases, the systematic error of MINFLUX scales with *L*^2^. Hence, reducing *L* effectively suppresses the positional bias and shifts its maximum towards smaller inclination angles producing much weaker fluorescence (Fig. 2b,c). Close to the excitation zero, the dipole-projected transverse field varies linearly with distance, whereas the longitudinal contribution increases quadratically. The corresponding excitation rate therefore contains a leading symmetric term that scales with the square of the distance from the probing intensity zero and a smaller asymmetric term that scales with its cube. Because the probe distance is proportional to *L*, the leading-order scaling is *L*^2^ sin^2^ *θ* for the symmetric signal and *L*^3^ sin *θ* cos *θ* for the asymmetric signal. Normalizing the asymmetric signal by the total signal gives a factor proportional to *L* |cot *θ*|.

Converting this result into a position introduces another factor of *L*, yielding *Δr* ∝ *L*^2^ |cot *θ*| (Fig. 2b,c). This leading-order relation dominates the behavior of the systematic error except for fluorophores with *θ* close to zero, which are largely perpendicular to the focal plane and hence dim. The maximum error scales with *L*, as has been reported earlier^29^. In MINFLUX, residual systematic errors due to orientational effects are therefore controllable by the recording parameters.

The fluorophores used in practical microscopy are attached to the biomolecules of interest, either directly or through a molecular linker. Attachment inevitably prevents a fluorophore from rotating freely and hence from assuming all orientations during photon collection. We have modeled this restricted mobility by allowing the fluorophore to sample orientations within a cone of half-angle *α* around a mean orientation (Fig. 2d). Increasing *α* averages over more emission patterns and therefore reduces the orientation-dependent camera bias. For a cone centered at an inclination of 45°, however, the bias remains almost 5 nm at *α* = 70° (Fig. 2e). Only in the unrealistic limit of complete rotational averaging is the orientation-dependent error fully removed. In contrast, throughout the same range, MINFLUX remains well below the error seen with camera-based localization. Therefore, the practically relevant question is how strongly attachment restricts rotation, because the degree of restriction determines the remaining accuracy error.

### An orientation ensemble defocus assay measures localization accuracy without positional ground truth

A fixed molecule does not reveal its true coordinate, and its dipole orientation is generally unknown. Yet, the accuracy becomes experimentally detectable by scanning a fixed-orientation fluorophore along the optical axis (Fig. 1c,d). To investigate the extent of the systematic error for all possible orientations both in theory and practice, we designed an experiment featuring an ensemble of many randomly oriented but fixed molecules residing in the same focal plane. We fixed the dipoles by spin-coating them into a polystyrene layer. As this layer is moved along the *z*-axis, each fluorophore follows a different lateral localization trajectory as *z* changes, depending on its fixed orientation.

Aligning the mean position of each trajectory removes its arbitrary position within the focal plane. Analyzing the width of the remaining distribution at each *z*-position isolates the orientation-dependent movement because errors that affect all molecules equally are cancelled out. Note that alignment by subtracting the mean neither favors the central focal point nor does it insert the assumption that the lateral position at *z* = 0 is without error. This ensemble experiment therefore allows unknown molecular orientations to be converted into a reproducible macroscopic distribution whose shape and width can be used to test and quantify systematic errors induced by defocus.

A simulation of the experiment showed that the axial trajectories of the camera localizations form a broad double cone around the focus (Fig. 3a). Their lateral spread increases with |*z*| because defocus turns the pupil phase of each emitting dipole into an orientation-specific displacement. Under a brightness threshold used to represent practical microscopy and post-processing conditions, the systematic spread 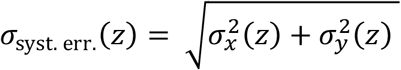 reaches 31.4 nm at *z* = ±300 nm (Fig. 3c).

**Fig 3.**
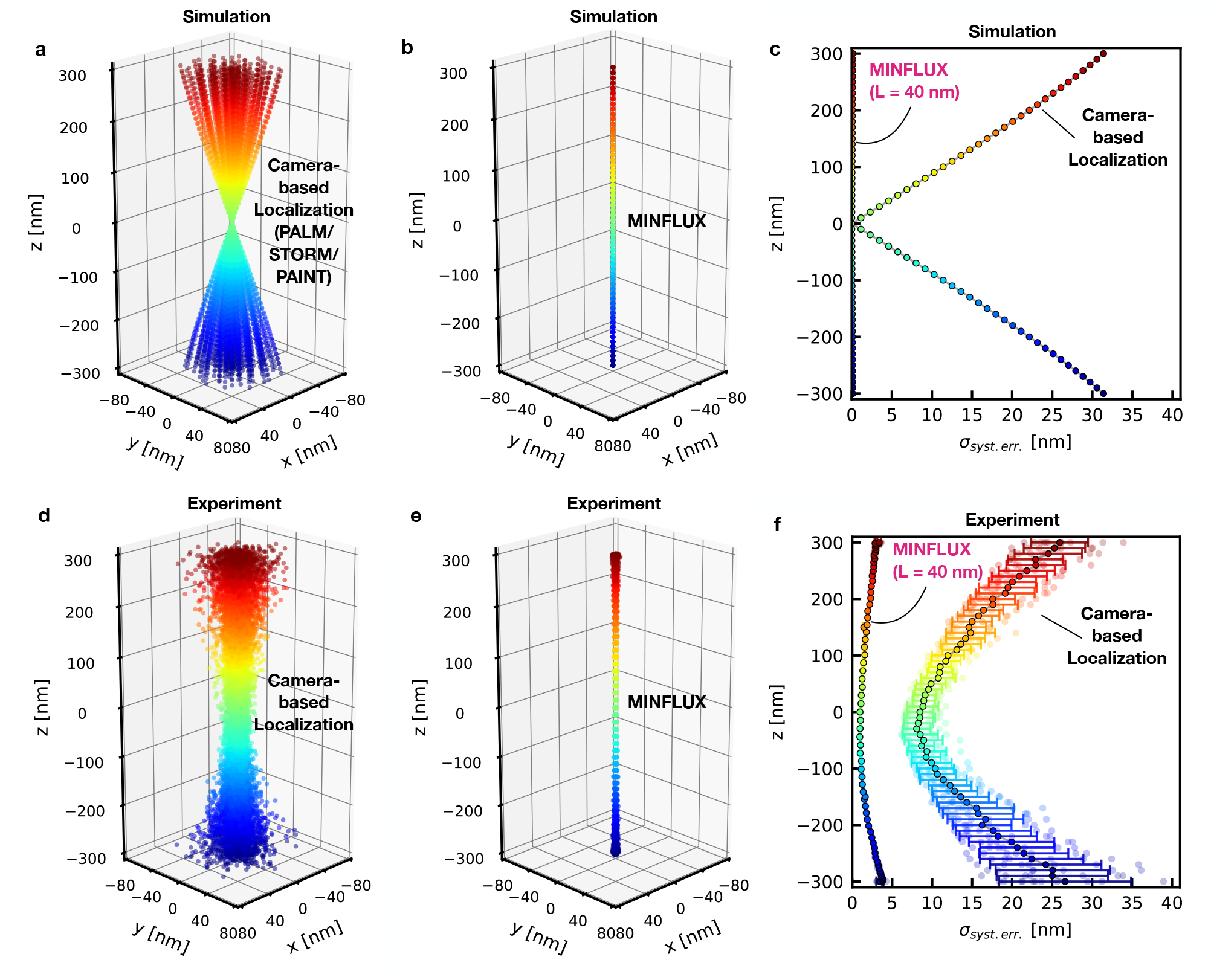
Simulated and measured systematic positional (accuracy) errors for fixed dipoles. **a**,**b**, Simulated orientation-dependent localization trajectories as a function of defocus z for fixed dipoles with uniformly sampled orientations, using camera-based localization (**a**) and MINFLUX with L = 40 nm (**b**). Simulation parameters and estimators as in Figs. 1 and 2, except for a centroid window of ±300 nm. Each trajectory was centered by subtracting its mean lateral position. Orientations below a realistic brightness threshold were excluded. **c**, Systematic positional spread, 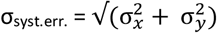, calculated from the standard deviations of the aligned positions across dipole orientations in x and y. The simulations contain n = 101 camera-localized and n = 106 MINFLUX-localized orientations after brightness selection. **d**,**e**, Aligned experimental trajectories from representative ensembles of fixed fluorophores (ATTO 647N-biotin) measured by camera-based localization (n = 237; **d**) and MINFLUX (n = 33; **e**) over z = −300 to +300 nm. Experimental camera localizations were obtained by maximum-likelihood fitting of an integrated Gaussian point-spread function; MINFLUX localizations were provided by the commercial instrument. **f**, Systematic positional spread for k = 12 camera and k = 6 MINFLUX measurements, each containing at least 30 fluorophores. Black-outlined markers show the mean across ensembles and error bars show the standard deviation.

Here, *σ*_*x*_(*z*) and *σ*_*y*_(*z*) are the standard deviations across the aligned molecule positions in the two lateral directions. If dim molecular orientations are also included, the value increases to 48.4 nm. Under the same conditions, the simulated MINFLUX trajectories remain concentrated around the imposed optical coordinate (Fig. 3b). For a final probing diameter of *L* = 40 nm, as used in the default acquisition sequence of our MINFLUX microscope, the corresponding spread is 0.54 nm when all orientations are included. If we then apply the same brightness-selection rule as for camera localization, the error spread shrinks to 0.13 nm, which is 233-fold smaller than the camera-localization spread.

In our camera-based localization experiment, axial defocus was introduced by translating the sample along the optical (*z*) axis, whereas in the MINFLUX setup, defocus was generated by imposing a phase modification on the illumination wavefront while maintaining alignment of the confocal pinhole. Applying a defocus phase modification allowed us to perform a fast all-optical axial sweep on a single fluorophore. We selected -300 nm < z < 300 nm since it largely corresponds to the axial full width at half maximum of the optical detection. The corresponding measurements yield the same geometry of the axial trajectories as in our simulations (Fig. 3d-f). For camera-based localization, a representative ensemble of 237 fixed fluorophores expands from a measured *σ*_syst.err._ = 6.79 nm at *z* = 0 to 22.65 nm at *z* = -300 nm and 21.51 nm at *z* = +300 nm (Fig. 3d). For MINFLUX localization with an identically prepared sample containing 33 fixed fluorophores, the spread remains confined to *σ*_syst.err._ = 1.13 nm at *z* = 0, 3.69 nm at *z* = -300 nm, and 3.15 nm at *z* = +300 nm (Fig. 3e). Across multiple independent ensembles, camera localization gives *σ*_syst.err._ = 8.5 ± 1.6 nm at *z* = 0 and 27 ± 8 nm and 26 ± 4 nm at *z* = -300 and +300 nm, respectively (Fig. 3f). The MINFLUX measurements show *σ*_syst.err._ = 0.97 ± 0.09 nm at *z* = 0 and 3.45 ± 0.44 nm at *z* = -300 nm and 3.38 ± 0.23 nm at *z* = +300 nm (Fig. 3f).

The measured camera localization curve decreases to a finite value of *σ*_syst.err._ = 8.5 ± 1.6 nm at *z* = 0 nm rather than to the theoretically expected zero. This is because the finite *N* of available photons limited our localization precision *σ*_2D_ to 7.8 ± 1.7 nm which inherently set a lower bound on the measurable camera-localization accuracy. In our camera-localization experiments, increasing *σ*_2D_ by increasing *N* was not an option, because the limited photon budget of a single fluorophore had to be partitioned among the different *z*-measurements. This partitioning is necessary to ensure that the same uniquely oriented fluorophore is measured across the whole z-range. Yet the attained *σ*_2D_ is sufficient to confirm that the underlying theory governs the practical accuracy and to estimate an accuracy improvement of MINFLUX over camera-based localization by about 8-fold.

For MINFLUX, the two-dimensional precision of each molecule’s mean position is about 16 times higher: *σ*_2D_ = 0.48 ± 0.23 nm. The higher precision allows us to draw more detailed conclusions for MINFLUX. For example, the slightly larger values of *σ*_syst.err._ in and out of focus cannot be due to localization precision alone, as they also were not significantly reduced when reducing *L*. This suggests that they arise due to aberrations, sample heterogeneity or any residual errors that are slightly different for each individual molecule. Indeed, when assuming commonly encountered amounts of spherical aberration of 0.1 *λ* and 0.1 *λ* astigmatism in our simulation, the expected position errors increased to 0.15 nm at *z* = 0 and 3.50 nm at *z* = +300 nm and 3.11 nm at *z* = -300 nm, matching our experimental results.

### Common fluorophore attachment geometries retain dipole-induced systematic errors

Superresolution imaging and localization applications in biology often treat attached fluorophores as freely rotating. To test this assumption, we investigated different fluorophore attachment geometries with different degrees of freedom (Fig. 4). It is virtually impossible to prepare a ‘freely’ rotating fluorophore because any linkage to a biomolecule is restrictive. However, a freely rotating fluorophore is readily mimicked by a 23 nm sized fluorescent bead containing many fluorophores. As these fluorophores are randomly oriented, their joint fluorescence emulates that of a freely rotating fluorophore averaging over all orientations in space. Indeed, scanning fluorescent beads along the optical axis confirms the theoretical expectation that the camera-based localization spread changes only weakly, from 6.5 ± 0.4 nm in focus to 8.3 ± 1.6 and 8.6 ± 1.6 nm at *z* = -300 and +300 nm (Fig. 4a). Due to its higher localization precision for a given *N*, localization by MINFLUX reduces the same spread to 0.73 ± 0.39 nm in focus and 2.0 ± 0.4 nm and 2.2 ± 0.6 nm at the defocus limits. In any case, the orientation-induced C-shaped dependence is essentially absent in both approaches, in line with our calculations that the systematic errors are due to a combination of defocus and orientation.

**Fig 4.**
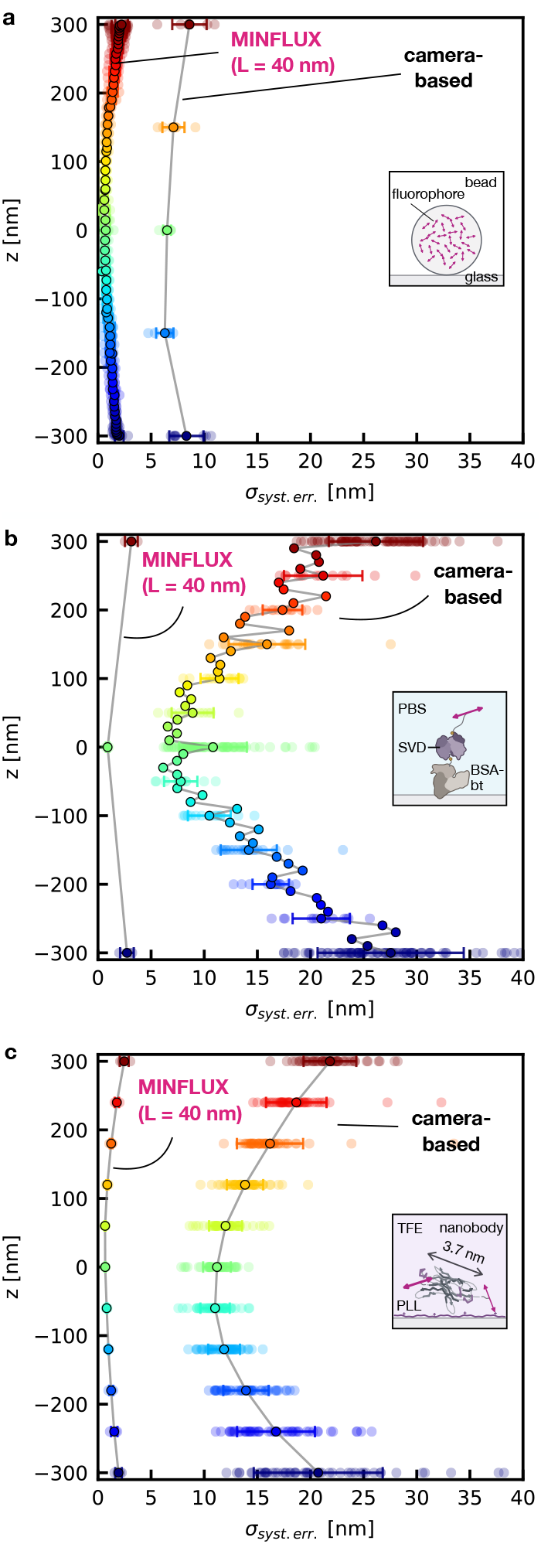
Systematic positional errors for standard samples with some degree of fluorophore rotational mobility. **a–c**, Systematic positional spread σ_syst.err._, defined in Fig. 3, as a function of defocus for camera-based localization and MINFLUX. Samples comprise fluorescent beads containing multiple randomly oriented fluorophores (**a**), single fluorophores (ATTO 647N-biotin) tethered through a BSA–biotin–streptavidin linkage (**b**) and fluorophores conjugated to surface-immobilized nanobodies (**c**). For the nanobody sample, each nanobody carried two fluorophores, of which one was uncaged during each measurement. Camera and MINFLUX localizations were obtained using the same localization procedures described in Fig. 3. Each ensemble contained at least 30 beads or fluorophores. Markers show the mean across k ensembles and error bars show the standard deviation. The numbers of ensembles for camera/MINFLUX were k = 8/8 in **a**, k = 86/3 in **b** and k = 50/4 in **c**.

Next, we carried out the same axial scanning with single fluorophores attached through a BSA-biotin-streptavidin linker embedded in phosphate-buffered saline (PBS). Here, we expected the fluorophore to enjoy substantial rotational freedom, but also restrictions due to the linker. Indeed, the result shows a substantial increase in systematic localization errors for camera localization (Fig. 4b), with the spread rising from 11 ± 3 nm in focus to 28 ± 7 nm at -300 nm and 26 ± 4 nm at +300 nm. The pronounced C-shaped curve proves that the fluorophore does not freely average over all orientations. Note that no independent measurement of rotational mobility is required to reach this conclusion, because free rotation would essentially remove the C-shape, as demonstrated by the bead control. Practical MINFLUX localizes these fluorophores with 0.91 ± 0.16 nm spread in focus and only 2.7 ± 0.6 nm and 3.1 ± 0.6 nm at the defocus limits, showing that it is by an order of magnitude more resilient against the systematic localization errors stemming from the rotational restriction by the linker.

Since fluorophores are widely used to label proteins directly, e.g. through a maleimide or amine chemistry, we also investigated the systematic errors associated with the localization of fluorophores bound to nanobodies. The nanobodies were spread out on a poly-L-lysine-coated glass coverslip and imaged in trifluoroethanol (TFE). The camera-based data again trace the C-shaped signature, increasing from 11.2 ± 1.3 nm in focus to 21 ± 6 and 21.8 ± 2.5 nm at -300 and +300 nm (Fig. 4c), revealing that this labeling method provides rather incomplete fluorophore rotation. In contrast, MINFLUX confines the spread to 1.9 ± 0.3 nm at -300 nm, 2.5 ± 0.4 nm at +300 nm and 0.68 ± 0.05 nm in focus (Fig. 4c).

Together, both experiments reveal that the magnitude of the accuracy error depends on the fluorophore attachment geometry. Yet, samples that are sometimes assumed to provide enough rotational freedom, in fact still entail a 10-15 nm deviation from ground truth that can be of the same order as for fixed dipoles.

### MINFLUX preserves nanometer-scale molecular distances under realistic defocus

The recent achievement of ångström localization precisions has spurred initial applications of superresolution fluorescence microscopy to structural biology, specifically to the measurement of single protein (complexes) and conformational states of individual biomolecules^14,17^. It goes without saying that a reliable application of superresolution in this field requires high levels of accuracy as much as it requires precision. In fact, most applications in this emerging field boil down to measuring distances between two or more fluorophores on different sites of a biomolecule. If the fluorophores do not have the same orientation, these measurements are prone to errors because any defocus changes the measured distance between them. Such errors are particularly detrimental when measuring distances, especially when the distances are small.

Using our MINFLUX setup, we localized two fluorophores attached to the N and C termini of the same nanobody. The fluorophores were known to be separated by 3.7 nm as already demonstrated by Sahl et al.^17^ using MINFLUX. Since defocus plays a critical role, we decided to settle on a defocus range that is difficult to detect by a decrease in brightness. At *z* = ±120 nm, the peak brightness of a fluorophore is still about 94% of its peak at *z* = 0, meaning that it is rather generous to assume that the -120 nm < z < 120 nm range can be tightly controlled without further precautions. For a nanobody with two fluorophores precisely localized with high accuracy, the distance changed by only 0.5 ± 0.6 nm (Fig. 5a). For a median-case example, the distance changed by 1.4 ± 0.4 nm over the full ±120 nm axial range (Fig. 5b). At *z* = 0, the distance distributions featured medians of 3.81 nm for *L* = 20 nm and 4.31 nm for *L* = 40 nm, close to the expected 3.7 nm distance (Fig. 5d,e). We note that the close agreement of the measured distance with the reported value of 3.7 nm is yet another proof of the accuracy provided by MINFLUX, as is essential for intra-molecular scale analyses^17^.

**Fig 5.**
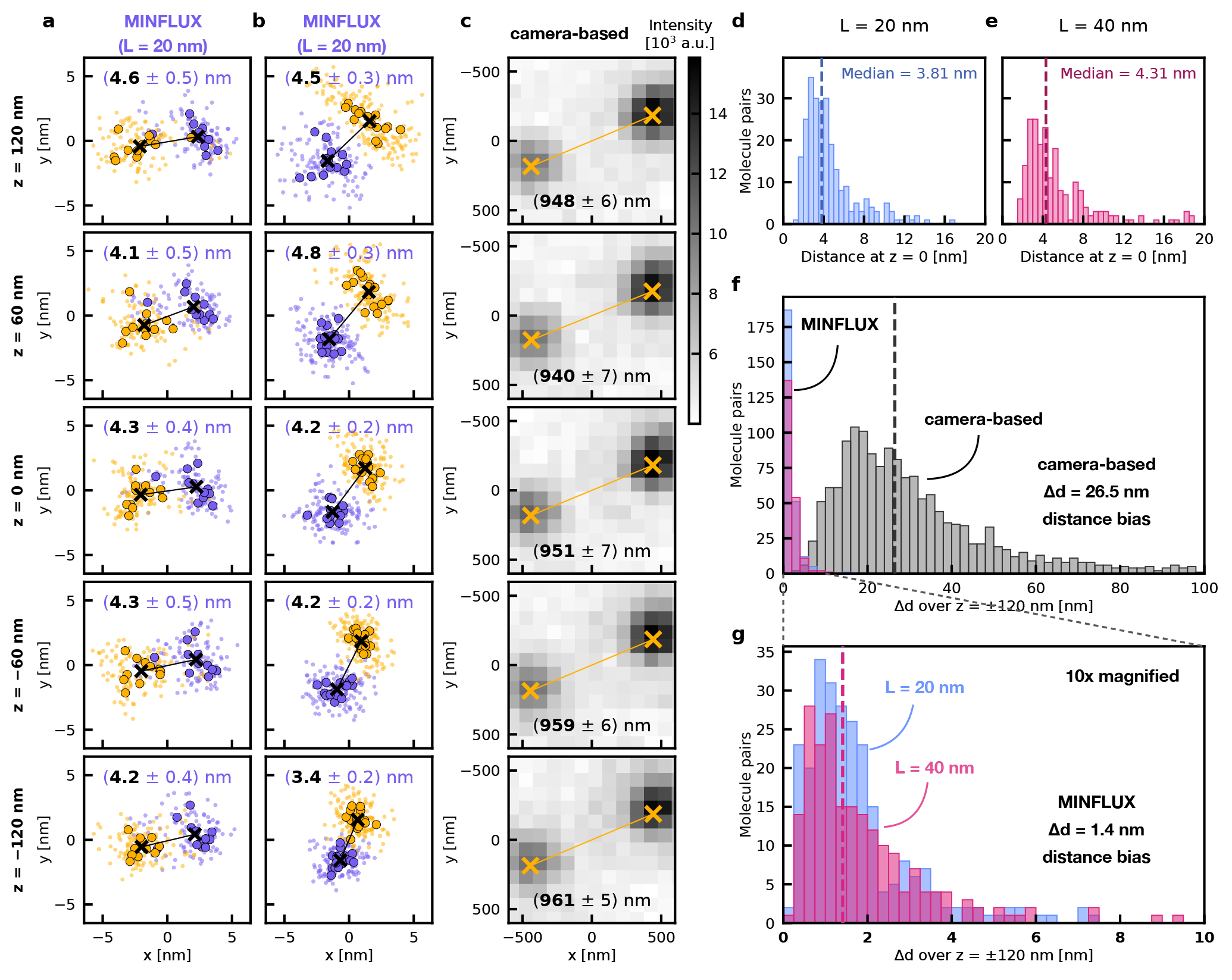
Defocus-dependent variation in molecular-distance measurements. **a**,**b**, MINFLUX measurements with L = 20 nm of the distance between fluorophores attached to the N and C termini of the same nanobody, whose attachment sites are separated by 3.7 nm. Two example traces show total distance changes of Δd = 0.5 ± 0.6 nm (**a**) and 1.4 ± 0.4 nm (**b**) over z = −120 to +120 nm. In **a**,**b**, localizamons were assigned to dye 1 (orange) and dye 2 (purple) using instrument-provided fluorescence trace idenmfiers (TIDs). Two TIDs were assigned to dye 1 and three to dye 2 in **a**, whereas one TID per dye was used in **b. c**, Example camera-based measurement of the separation between fluorophores on different nanobodies, with Δd = 21 ± 9 nm over the same range. Here, Δd denotes the difference between the maximum and minimum measured distance within the axial interval. **d**,**e**, In-focus distance distributions measured by MINFLUX with L = 20 nm (n = 256 molecule pairs; **d**) and L = 40 nm (n = 208 molecule pairs; **e**). The respective medians are 3.81 nm and 4.31 nm. **f**, Distributions of Δd for camera-based localization (n = 1,550 pairs) and MINFLUX (n = 256 pairs at L = 20 nm and n = 208 pairs at L = 40 nm). **g**, Enlarged view of the MINFLUX distribution in **f**. Median Δd values are 26.5 ± 0.8 nm for camera localization and 1.41 ± 0.06 nm for MINFLUX.

With camera-based localization, however, the measured distance between two fluorophores on different nanobodies changed in a typical example by 21 ± 9 nm across the same defocus range (Fig. 5c). Across 1,550 camera-localized pairs, the median change was 26.5 ± 0.8 nm (Fig. 5f). This error exceeds the complete dimension of many protein complexes and clearly overwhelms a 3.7 nm molecular separation. In stark contrast, the median defocus-induced change across 463 distances measured by MINFLUX was 1.41 ± 0.06 nm (Fig. 5f,g).

## Discussion and Outlook

We have shown that the systematic localization errors of individual fluorophores arising from their inherently unknown orientation and axial defocus compromise the straightforward applicability of camera-based localization to the sub-10 nm scale. In practical MINFLUX localization, in its current state of development, such systematic errors are reduced by about an order of magnitude. In theory, these errors can be virtually absent. We found that for rotationally restricted fluorophores, a commercial MINFLUX microscope with potential for further improvement delivers a roughly 8 times higher accuracy than camera-based localization.

Whereas the accuracy limits of centroid-based camera localization are grounded in the physical laws of fluorescence emission, the attained MINFLUX accuracies are not. In fact, both our calculations and experiments indicate that the residual errors of MINFLUX can be reduced to a fraction of an ångström. An obvious route towards this goal is to take advantage of the scaling of the accuracy value with *L*^2^. For example, at *θ* = 45°, reducing the probing diameter *L* from the typically used 40 nm to 10 nm decreases the calculated error from 8.5 Å to 0.53 Å, i.e., by 16-fold. The same reduction of *L* improves precision only fourfold. On the other hand, the corresponding experiments (Fig. 5g) indicate that reaching sub-ångström accuracies is challenged by residual aberrations of the doughnut-shaped illumination beam, indicating that higher levels of accuracy call for active aberration compensation. The fact that our results were obtained from a commercially available MINFLUX system and not from a perfected laboratory instrument highlights their practical relevance. At the same time, they underscore that optimizations will lead to additional improvements.

A route to attaining similar accuracy in camera-based localization would require a tight control of the axial coordinate *z* = 0, for example by additional optical control loops. However, in a three-dimensional sample, such as a cell, sub-50 nm controls of the defocus are difficult to realize if the fluorophores are not confined to a single plane. It is often assumed that the rotational mobility of fluorophores is sufficient to eliminate systematic positional errors altogether. However, with the common fluorophore attachment geometries investigated here, the observed spreads in camera-based localization remained surprisingly comparable to those of fixed fluorophores. Thus, in camera-based localization, attachment to a protein or a linker does not guarantee sufficient rotational freedom for systematic errors to average out. Instead, our theoretical study shows that even a generous full cone angle of 140° does not allow for enough rotational averaging if the fluorophore is tilted relative to the focal plane by 45° on average.

While it is generally accepted that future fluorophore linkers should become smaller in order to minimize the offset between the biomolecule and the fluorophore, our work shows that, for camera-based localization microscopy, developing linkers that provide high rotational freedom is at least as important. If the linkers do provide flexibility, it is important that the fluorophore makes use of it and does not stick to surrounding biomolecules, which may have been the case in our experiments using the fluorophore Atto647N as a label. Of equal relevance is the embedding medium, which co-determines the orientational space covered by the fluorophore when delivering the *N* photons for an individual localization. Clearly, the larger *N* is, the higher the probability that orientational effects average out to the degree allowed by the cone angle *α* for a certain mean orientation *θ*. Therefore, methods requiring *N* ≫10,000 photons per fluorophore, or even fluorophore replenishment for the same localization, such as DNA-PAINT^15^, may eventually lead to smaller accuracy errors. Also, the accuracy error is less relevant for distance measurements if the fluorophores at the endpoints have virtually identical spatial orientation and the same *z*-coordinate. While such exceptional situations are likely to occur in certain biomolecular arrays and two-dimensional binding templates, this cannot be the default situation in a molecular complex and in a cell. For MINFLUX, our work shows that these aspects of molecular orientation are by far less relevant.

It is also worth noting that the actual consequence of a given fluorophore tilt is a specific, elaborate deformation of its fluorescence spot on the camera. In principle, one could reconstruct the magnitude of the tilt^25,35,36^, albeit with an orientational ambiguity, and recover the correct position if the number *N* of detected photons were sufficiently high to permit perfect fitting of this deformation^36^. Localization microscopy resorts to generic centroid calculation or fitting with a Gaussian because the number *N* of photons provided by the fluorophore is by far lower. While each of these evaluation methods has its advantages in particular settings, none of them can mitigate the accuracy problem. Resorting to a different evaluation modality in our study would not have changed its central outcome. This reasoning also highlights the conceptual advantage of localization by MINFLUX: the burden of requiring many photons to obtain a coordinate-defining photon pattern, as inherently demanded by Heisenberg’s uncertainty principle, is taken from the fluorophore and placed on the illumination beam, where the number of photons is virtually unlimited and their polarization is readily controlled. Although it remains to be shown, the same conceptual argument applies to MINSTED^10,11,37^, a related coordinate-targeted localization method that also provides ångström precision.

In a nutshell, attaining (sub)nanometer localization precision is not sufficient to extend superresolution fluorescence microscopy or nanoscopy into the sub-10 nm domain. By providing both the precision and the required accuracy, coordinate-targeted localization, as shown here in the form of MINFLUX, has all the prerequisites for developing optical methods that investigate the dimensions and conformations of individual biomolecules reliably.

## Acknowledgements

We thank our colleague Marcel Leutenegger for providing MATLAB code used to double-check the calculations and for advice and discussions regarding simulations and data analysis. We are grateful to our colleague Steffen Sahl for valuable discussions regarding the use of nanobodies for distance measurements and preparing nanobody samples. We thank both for critically reading the manuscript. We thank Abberior Instruments GmbH for providing access to one of its MINFLUX systems. K.B.C. is a member of the Max Planck School of Photonics.

## Author contributions

S.W.H. proposed and supervised the project and is responsible for its scientific thrust direction. K.B.C. implemented the experimental approach for quantifying accuracy, prepared the samples, performed the experiments, simulations and data analysis in continual feedback with J.E. and S.W.H. R.S. adapted the localization software of the used commercial MINFLUX microscope to enable synchronized axial scanning. J.E. implemented synchronized axial scanning on the camera-based localization microscope. All authors discussed the results and implications of this study. The paper was written by K.B.C. and S.W.H.

